# Genetic diversity and population structure of *Luffa acutangula* accessions in a Thailand collection using SNP markers

**DOI:** 10.1101/2020.07.16.206128

**Authors:** Grimar Abdiel Perez, Pumipat Tongyoo, Julapark Chunwongse, Hans de Jong, Paweena Chuenwarin

## Abstract

This study explored a germplasm consisting of 112 *Luffa acutangula* (ridge gourd) accessions mainly from Thailand, and some accessions from Vietnam, China, Philippines, Indonesia, USA, Bangladesh and Laos for an analysis of the population structure and underlying genetic diversity using 2,834 SNPs. STRUCTURE analysis (*ΔK* at *K*=6) allowed us to group the accessions into six subpopulations that corresponded well with the unrooted phylogenetic tree and principal coordinate analyses. The phylogenetic tree showed the diversity of *L. acutangula* in Thailand, and accessions from other countries apart from Thailand were grouped together in the same branches. In STRUCTURE, subpopulation 2 contained only accessions from Thailand while other subpopulations contained a combination of accessions from Thailand and from other countries. When plotted, the STRUCTURE bars to the area of collection, it revealed the geneflow from the surrounding places nearby as indicated by the admixed genetic in the STRUCTURE bars. AMOVA based on STRUCTURE clustering showed the variation between populations (12.83%) and confirmed the absence of population structure in subpopulations (−10.59%). There was a distinguishing characteristic fruit shape and length in each subpopulation. The ample genetic diversity found in the *L. acutangula* germplasm can be utilized in ridge gourd breeding programs to help meet the demands and needs of both consumers and farmers.

## Introduction

*Luffa acutangula*, commonly known as ridge gourd, angled loofah, or Chinese okra, is a domesticated vegetable of the Cucurbitaceae that originated from India (Filipowicz et al. 2014; Heiser and Schilling 1988; Pessarakli 2016; Soladoye and Adebisi 2004). Immature fruits are used as a vegetable which can be cooked or fried. In South-East Asia, the sweet juiciness, spongy texture, and mild-bitter taste are favorite characteristics (Pessarakli 2016). In contrast, its use in traditional medicine is especially prevalent in Asia and middle America (Morton 1981; Sastri 1962) and India for its treatment of jaundice and urinary bladder stones (Katewa et al. 2004; Samvatsar and Diwanji 2000). Extensive regional and cultural selection of specific cultivars have contributed to a wide variety of uses and value necessitating germplasm conservation to capture its enormous genetic variation. Most of the genetic diversity was established by describing the morphological variation of the accessions (Prakash et al. 2013). Still, such a limited approach is subject to environmental conditions as well as the developmental stages of the plant (Zhang et al. 2008). Instead, molecular markers provide more stable sources of information for carrying out genetic diversity studies.

Genetic diversity can be described as the measure of variation within a population. This variability is essential for future breeding schemes and represents the equilibrium between mutation and the loss of genetic variation (Leffler et al. 2012). The importance of population structure extends far beyond pure relatedness, in which plants with limited distribution have fewer habitats and smaller dense populations, which will lead to increased inbreeding and more structured populations (Holsinger 1991). If this structure is not correctly defined, it can lead to misinterpretation of genetic in the population of study. It is good to know if results using a particular method reflects similar findings in other methods of genetic assessment. The complementary assessments of genetic diversity which also serves to better define the relatedness from population structure are by the use of phylogenetic modeling and finding principal coordinate analysis (PCoA). These modelling helps to give an overview of the relative relatedness of each member of the population under study (Mardis 2008).

With the fast-paced innovations in molecular biology, several marker techniques have been developed for acquiring accurate and reliable information on population structure and genetic diversity of germplasms (Bretting and Widrlechner 1995). Molecular markers used for population structure and genetic diversity studies in ridge gourd include Inter Simple Sequence Repeat (ISSR) (Prakash et al. 2014), Random Amplified Polymorphism Detection (RAPD) in *L. acutangula* (Hoque and Rabbani 2009), Simple Sequence Repeats (SSRs) (Pandey et al. 2018), Inter Simple Sequence Repeats (ISSR), and Directed Amplification of Minisatellite DNA (DAMD) (Misra et al. 2017). These techniques can be labor-intensive, costly and produce a low number of markers.

The present study aims to identify the population structure and genetic diversity of a *L. acutangula* germplasm in Thailand by using DArTseq based SNPs (Kilian et al. 2012; Raman et al. 2014). The study will provide essential information for genetic improvement programs in Thailand for meeting the demands of a diverse cultural cuisine in Thailand.

## Materials and methods

### Plant materials

We used in this study a *L. acutangula* germplasm comprising of 112 accessions from Thailand (91), Vietnam (4), Philippines (2), Indonesia (1), China (3), USA (1), Laos (3) and Bangladesh (7), conserved by the Tropical Vegetable Research Center (TVRC), Kasetsart University, Kamphaeng Saen Campus, and the World Vegetable Center, Taiwan (**Online Resource Online** Resource S1).

### Fruit trait evaluation

Accessions were grown under field conditions from August to December 2019 in which five plants per plot were planted in a single bed with a 0.5 m space between the plants and 2 m spacing between beds. Each accession was self-pollinated. Fruits were harvested when they were young for consumption as vegetable. All *L. acutangula* accessions were evaluated for fruit traits such as fruit length by measuring the stem-end to blossom-end of three fruits per accession and fruit shape was adapted from the descriptors for sponge gourd (Joshi et al. 2004). Fruit shape was classified into either elongated slim, elliptical tapered, elongated tapered, elongated elliptical tapered, tapered oblong or short elliptical tapered.

### DNA extraction

Genomic DNA samples were extracted from 100 mg of pooled young leaves tissue of 2-weeks-old seedlings from 20 plants per accession using a modified CTAB (cetyltrimethylammonium bromide) method (Doyle and Doyle 1987). Precipitated DNA was resuspended in TE buffer (10 mM Tris-HCl; 1 mM EDTA, pH 8.0) containing 2 μg/mL RNase. DNA quality was evaluated by electrophoresis on a 1% agarose gel and was quantified with a NanoDrop 2000c spectrophotometer V 1.6.0. The DNA concentration was adjusted to 50 ng/μL for DArTseq GBS analysis.

### Genotyping of accessions of *L. acutangula* using DArTseq

The genomic DNA samples were sent to Diversity Arrays Technology Pty. Ltd., Canberra, Australia, for DArTseq genotype-based sequencing (Von Mark et al. 2013). To this end, DNA was digested using *Pst*I-*Mse*I restriction enzymes as described by Kilian (Kilian et al. 2012). The digested fragments were then ligated to adapters and amplified by PCR (Raman et al. 2014), followed by sequencing on Illumina Hiseq2000. The single read sequencing was run for 77 cycles, and sequences generated were handled by DArT analytical pipelines (Diversity Arrays Technology, Australia). In the primary pipeline, poor-quality sequences were filtered from the FASTQ files by applying rigorous selection criteria to the barcode region (Barilli et al. 2018). Identified sequences per barcode/sample were used for marker calling. These files were then used in the secondary pipeline for DArT P/L’s proprietary SNP calling algorithms (DArTsoftseq).

### Population structure and data analysis

DArTseq based SNPs were filtered using a call rate of 80% with a co-dominant marker polymorphism information content (PIC) greater than 0.125. After filtering, 2,834 SNPs were used for data analysis. The population structure of the 112 *L. acutangula* accessions was determined using STRUCTURE version 2.3.4 (Pritchard et al. 2000). Ten repeats were performed for each number of hypothetical subpopulations (*K*) which were set from 1 to 10. The parameters used consisted of an admixture model, a burning period of 50,000 steps, and 100,000 Markov Chain Monte Carlo (MCMC). The STRUCTURE results were further analyzed using the r package, pophelper version 2.3.0 (Francis 2017). The optimum number of *K* was calculated using the Evanno method (Evanno et al. 2005).

We constructed the phylogenetic trees with the weighted neighbor-joining method (Bruno et al. 2000) and visualized the data with DARwin software version 6.0.021 (Perrier and Jacquemoud-Collet 2006). Principal Coordinates Analysis (PCoA), expected heterozygosity (*H_E_*), observed heterozygosity (*H_O_*), and pairwise *F*_ST_ were calculated using adegenet version 2.1.1 (Jombart and Ahmed 2011) in the R statistical environment. Analysis of Molecular Variance (AMOVA) and Shannon-Weiner Diversity index were calculated in POPPR version 2.8.3 in R (Kamvar et al. 2014). Jaccard distance was calculated in r software by using ade4 version 1.7-13 package (Dray and Dufour 2007). One-way ANOVA test, box plots and Tukey HSD were carried out using standard r statistics.

## Results

### Population structure analysis

STRUCTURE analysis of the 2,834 SNPs revealed the highest value of *ΔK* at *K*=6 (**Fig. 1**). This value indicated a total of six informative subpopulations found across all *L. acutangula* accessions. Each subpopulation was expressed as Cluster 1-6, representing 7%, 55%, 13%, 8%, 4% and 13% of the total number of accessions, respectively. Cluster 1 consisted of seven accessions from Thailand and one from China (**Fig. 2**). All 62 accessions in Cluster 2 were from Thailand. Cluster 3 was composed of eleven accessions from Thailand and three from Laos. Cluster 4 was made of two, four, two and one accession from Thailand, Vietnam, Philippines and Indonesia, respectively. Cluster 5 consisted of two accessions from Thailand, two from China and one from the USA. Cluster 6 contained seven accessions from Thailand and seven from Bangladesh.

**Fig. 1.**
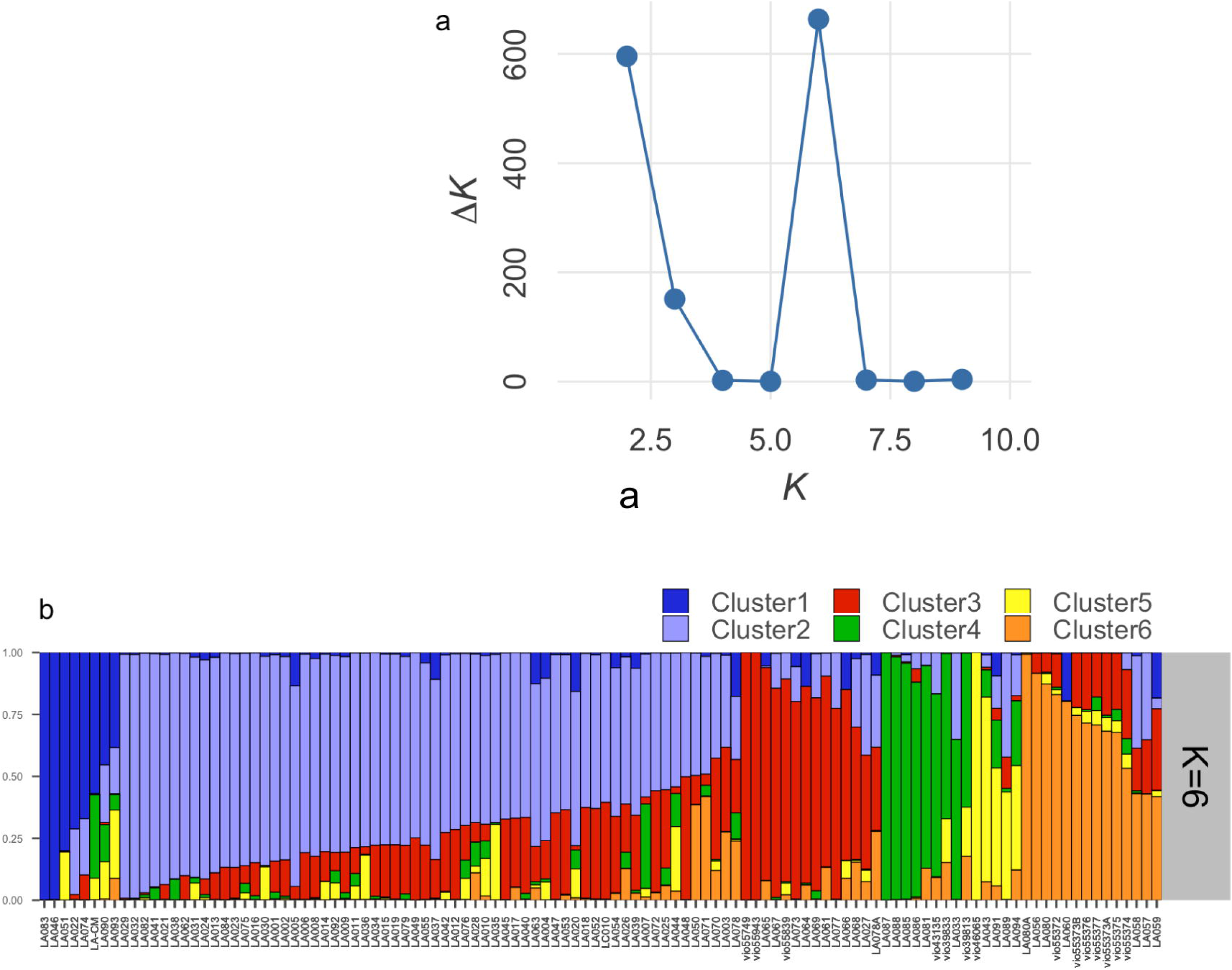
Population structure of 112 *Luffa acutangula* accessions using DArTseq-based SNP markers inferred by STRUCTURE program. **a** Number of subpopulations indicated by the highest *ΔK*; **b** Proportion of clustering of individuals to six subpopulations

**Fig. 2.**
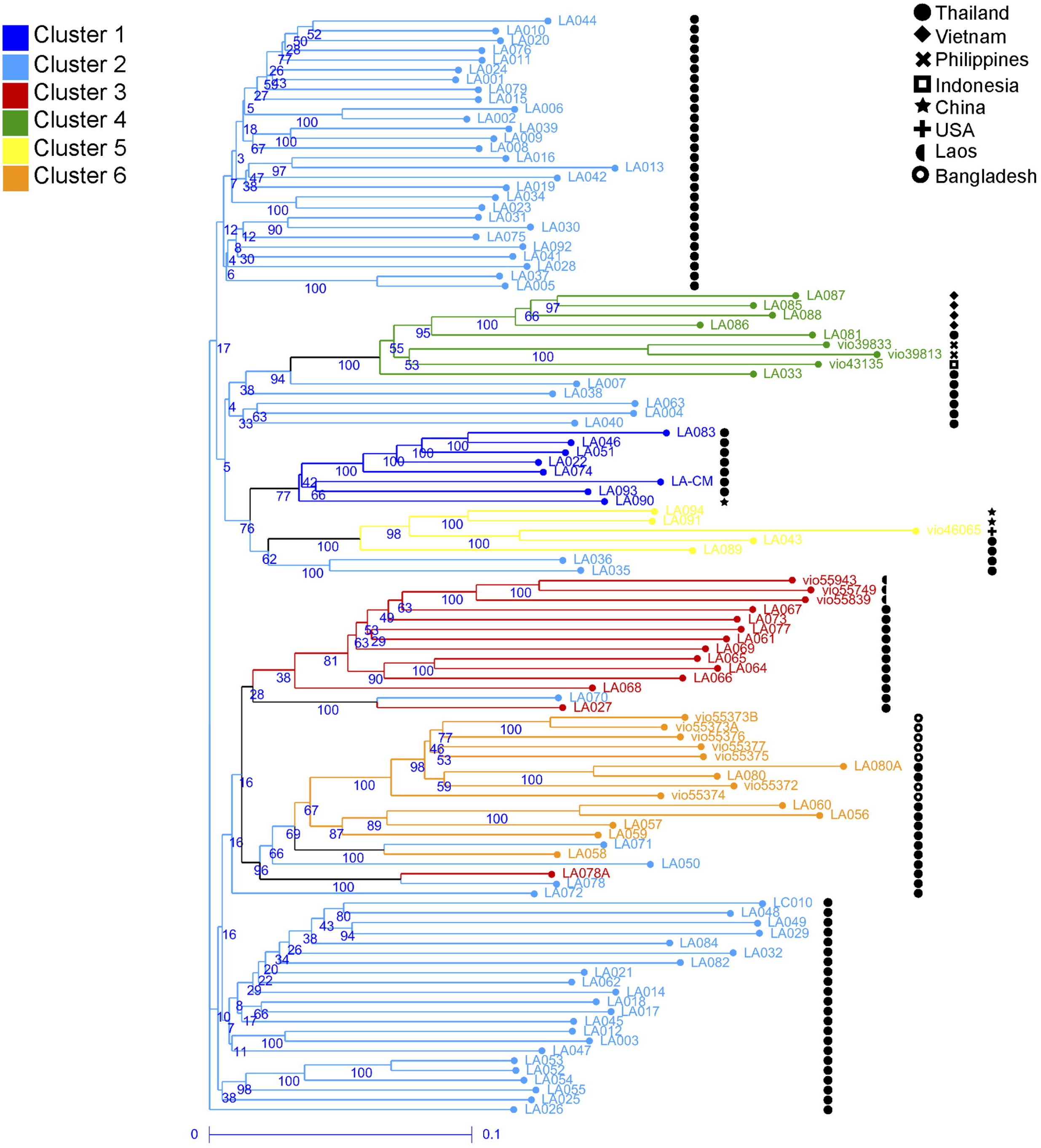
Weighted neighbor-joining dendrogram of 112 *Luffa acutangula* accessions based on DArTseq SNPs. Branch colors correspond to clustering by STRUCTURE analysis. Symbols indicate accession’s country of origin

The unrooted phylogenetic tree was able to differentiate the 112 *L. acutangula* accessions into six clades that were consistent with the *ΔK* at *K*=6 (**Fig. 3**). We further confirmed the distinction of the six groups with the Principal Coordinate Analysis (PCoA) in which the accessions of cluster 1 overlapped with those of cluster 2 (**Fig. 4**). Axes 1 and 2 combined explained 15.3% of the total variance. Cluster 1 and 2 had some accessions falling in two or three quadrants suggesting a wide diversity in these two clusters, which were mostly composed of Thailand accessions.

**Fig. 3.**
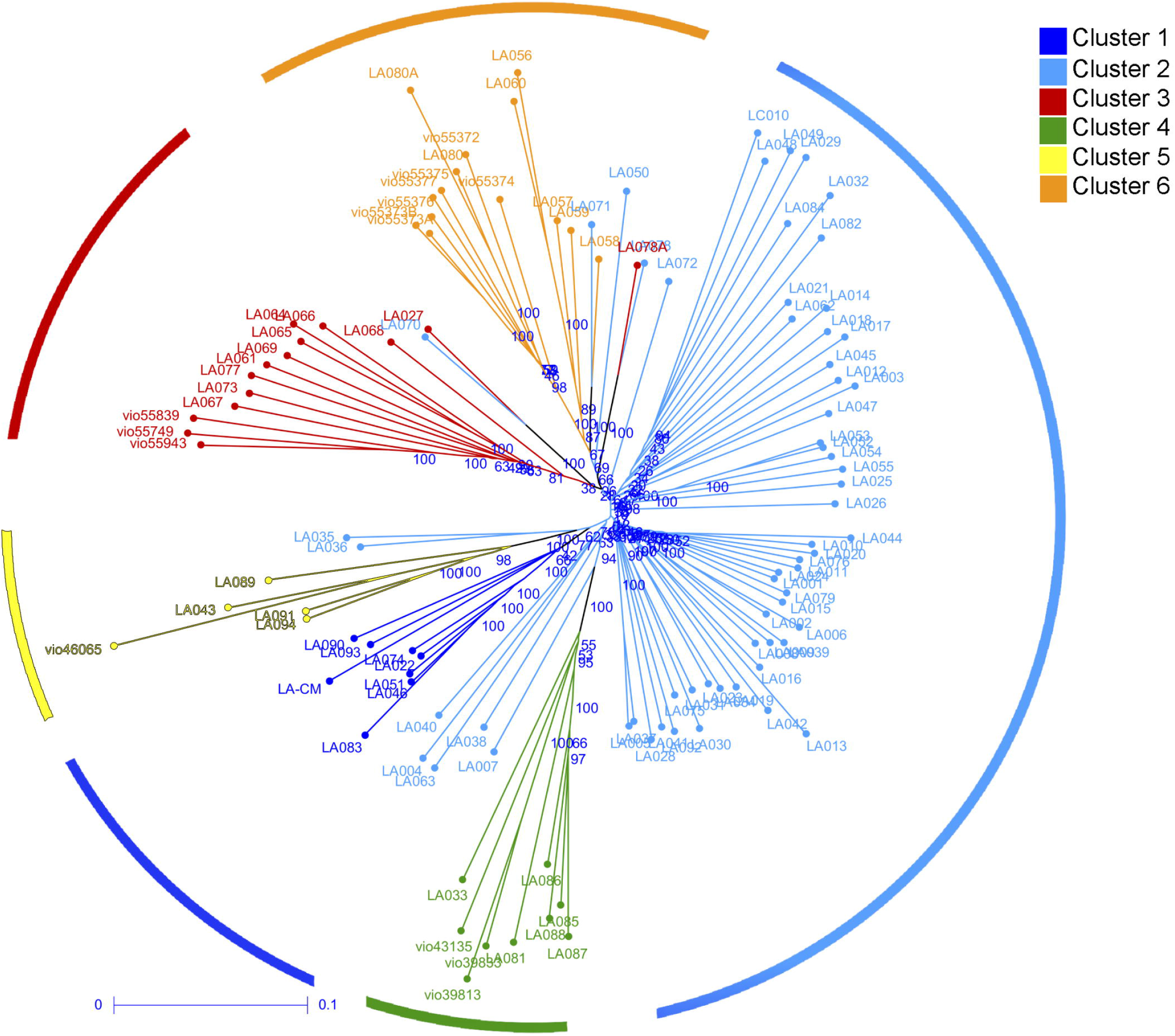
Phylogenetic tree among the 112 *Luffa acutangula* accessions. The colors of branches illustrate accessions belonging to different clusters acquired from STRUCTURE analysis. Six clades were identified as indicated by circle colors

**Fig. 4.**
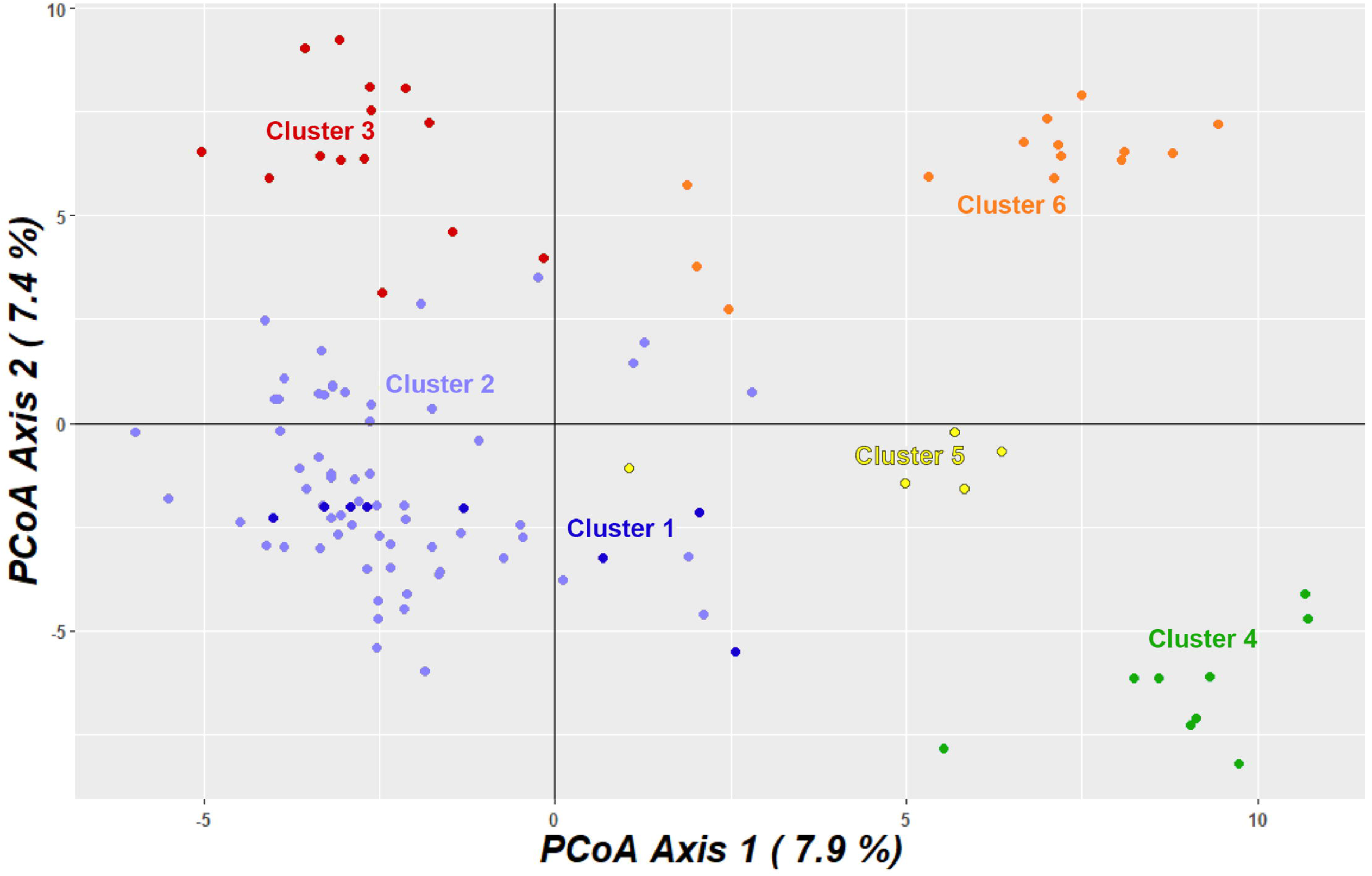
Principal coordinate analysis (PCoA) of 112 *Luffa acutangula* accessions using DArTseq-based SNPs

### Genetic diversity

Heterozygosity can be measured by expected heterozygosity (*H_E_*) and observed heterozygosity (*H_O_*). *H_E_* gives information about the probability of an individual’s portion of heterozygosity for all analyzed loci, while *H_O_* is the portion of heterozygous genes in the analyzed population (Chesnokov and Artemyeva 2015). The expected heterozygosity (*H_E_*) as measure of genetic diversity ranged in the six clusters from 0.338 (Cluster 2) to 0.272 (Cluster 3); the remaining cluster had values in between: Cluster 1 (0.274), Cluster 4 (0.295), Cluster 5 (0.287) and cluster 6 (0.334). The high *H_E_* in Cluster 2 also corresponded with the highest Shannon & Weiner diversity index (H) of 4.127. The observed heterozygosity (*H_O_*)for the STRUCTURE grouping was as follows: Cluster 1 (0.378), Cluster 2 (0.388), Cluster 3 (0.191), Cluster 4 (0.289), Cluster 5 (0.363) and Cluster 6 (0.410). When the observed heterozygosity was found higher than the expected heterozygosity, a random mating system governs that particular population (Chesnokov and Artemyeva 2015). This *H_O_* >*H_E_* was observed in Cluster 1, Cluster 2, Cluster 5 and Cluster 6 (Table 1). The observed heterozygosity (*H_O_*) in Cluster 4 (0.289) was almost similar in value to the *H_E_* (0.295), which might indicate that crossing in the population was almost unplanned (Chesnokov and Artemyeva 2015). Cluster 3 had the lowest *H_E_* (0.272) and *H_O_* (0.191) from all six subpopulations, and given that *H_O_* was lower than *H_E_* indicated that Cluster 3 was potentially an inbred population (Chesnokov and Artemyeva 2015). The mean SNP polymorphism information content (PIC) had a range value from 0.125 to 0.375, with a mean of 0.288 (Table 1).

**Table 1.**
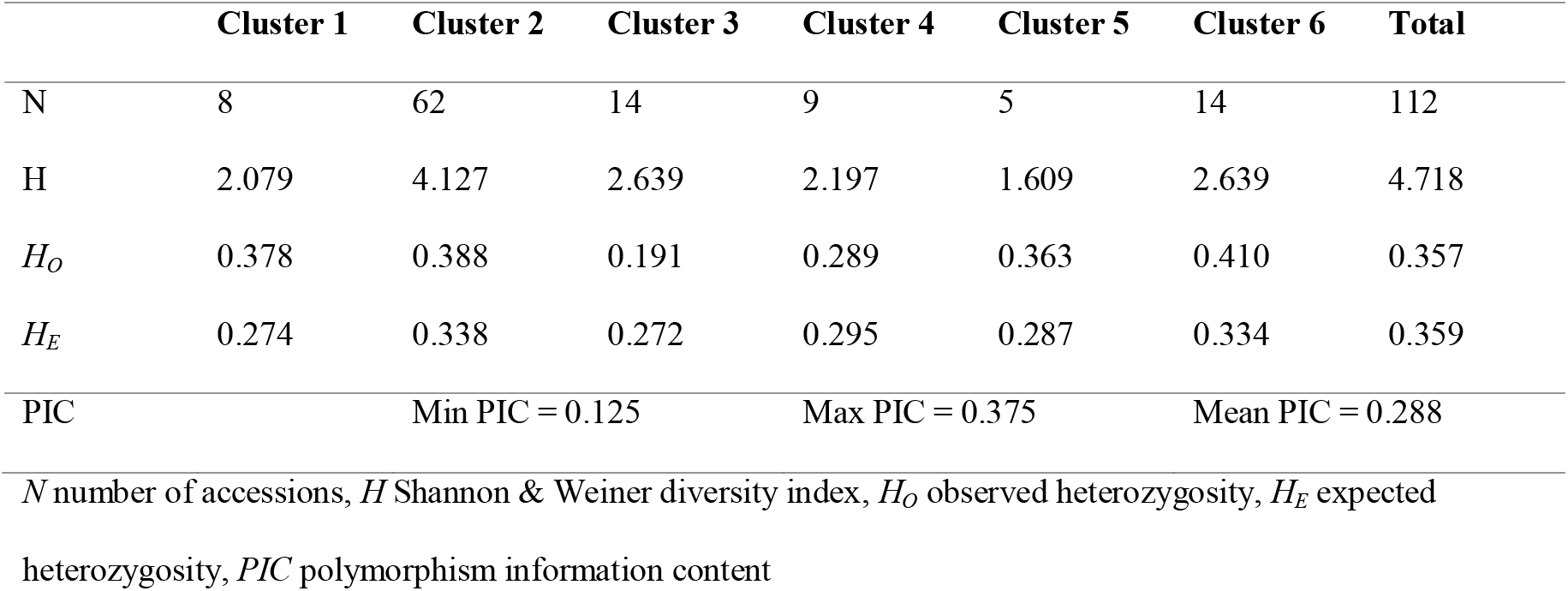
Genetic diversity among 112 *Luffa acutangula* accessions based on STRUCTURE analysis

Pairwise genetic differentiation was highest between Cluster 3 and Cluster 4 (*F*_ST_=0.262), while the lowest was observed between Cluster 1 and Cluster 2 (*F*_ST_ = 0.089), and also between Cluster 2 and Cluster 3 (*F*_ST_ = 0.086) (Table 2). AMOVA based on STRUCTURE clustering showed the 12.83% variation among subpopulations while negative AMOVA variance within subpopulations in STRUCTURE clustering (−10.59%) indicate the absence of genetic structure and emphasizes the nature of the outcrossing *L. acutangula* in which genes from different clusters can be more associated than genes from the very same cluster (Excoffier and Lischer 2009; Misra et al. 2017) (Table 3). Phi statistics between STRUCTURE clusters (0.128) indicate a moderate degree of differentiation (Frankham et al. 2002; Xu et al. 2019).

**Table 2.** Pairwise *F*_ST_ (genetic differentiation) values among clusters identified by STRUCTURE analysis

| STRUCTUR | Cluster 2 | Cluster 1 | Cluster 3 | Cluster 4 | Cluster 5 | Cluster 6 |
| --- | --- | --- | --- | --- | --- | --- |
| **E Clustering** |  |  |  |  |  |  |
| Cluster 2 |  |  |  |  |  |  |
| Cluster 1 | 0.089 |  |  |  |  |  |
| Cluster 3 | 0.086 | 0.180 |  |  |  |  |
| Cluster 4 | 0.164 | 0.230 | 0.262 |  |  |  |
| Cluster 5 | 0.147 | 0.203 | 0.202 | 0.210 |  |  |
| Cluster 6 | 0.123 | 0.186 | 0.127 | 0.193 | 0.174 |  |

**Table 3.**
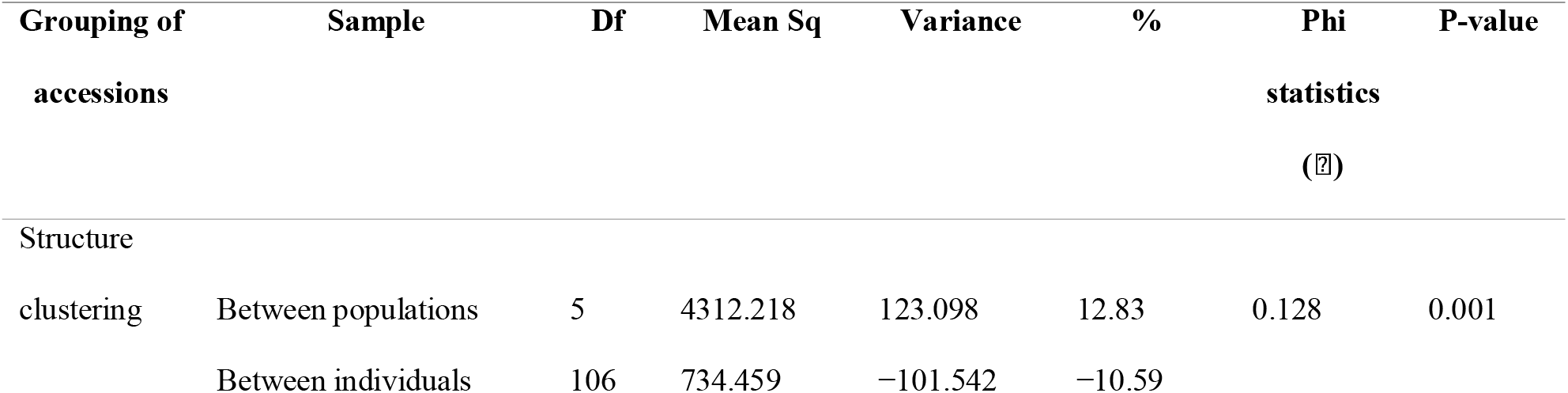
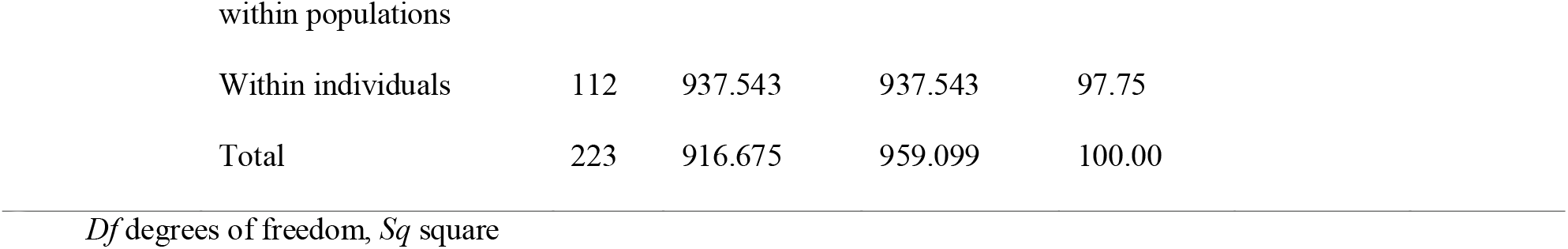
Analysis of molecular variance (AMOVA) of 112 *Luffa acutangula* accessions

### Geographic distribution and genetic diversity

Cluster 1 and 2 represented the genotype characteristics found in Thailand, while Cluster 3 consisted of characteristics belonging to Laos as is clear from the STRUCTURE analysis (**Fig. 5**). Accessions from Vietnam, Philippines and Indonesia were closely related with the majority of the shared genotypic characteristics from Vietnam in Cluster 4. Defining characteristics for Cluster 5 were from the USA, while for Cluster 6 were accessions from either Bangladesh or Thailand. In this study, the *L. acutangula* accessions from Thailand showed diversity that included the genetic proportions of *L. acutangula* accessions from other countries (**Fig. 6**). However, the accessions from Laos and Vietnam were highly uniform and had low genotype proportions from accessions belonging to other countries. Both the Philippines and Indonesia had genotypic proportions belonging to accessions from Vietnam and Bangladesh. Still, they differed for the Philippines which contained genotypes from the USA, while Indonesia had from Thailand.

**Fig. 5.**
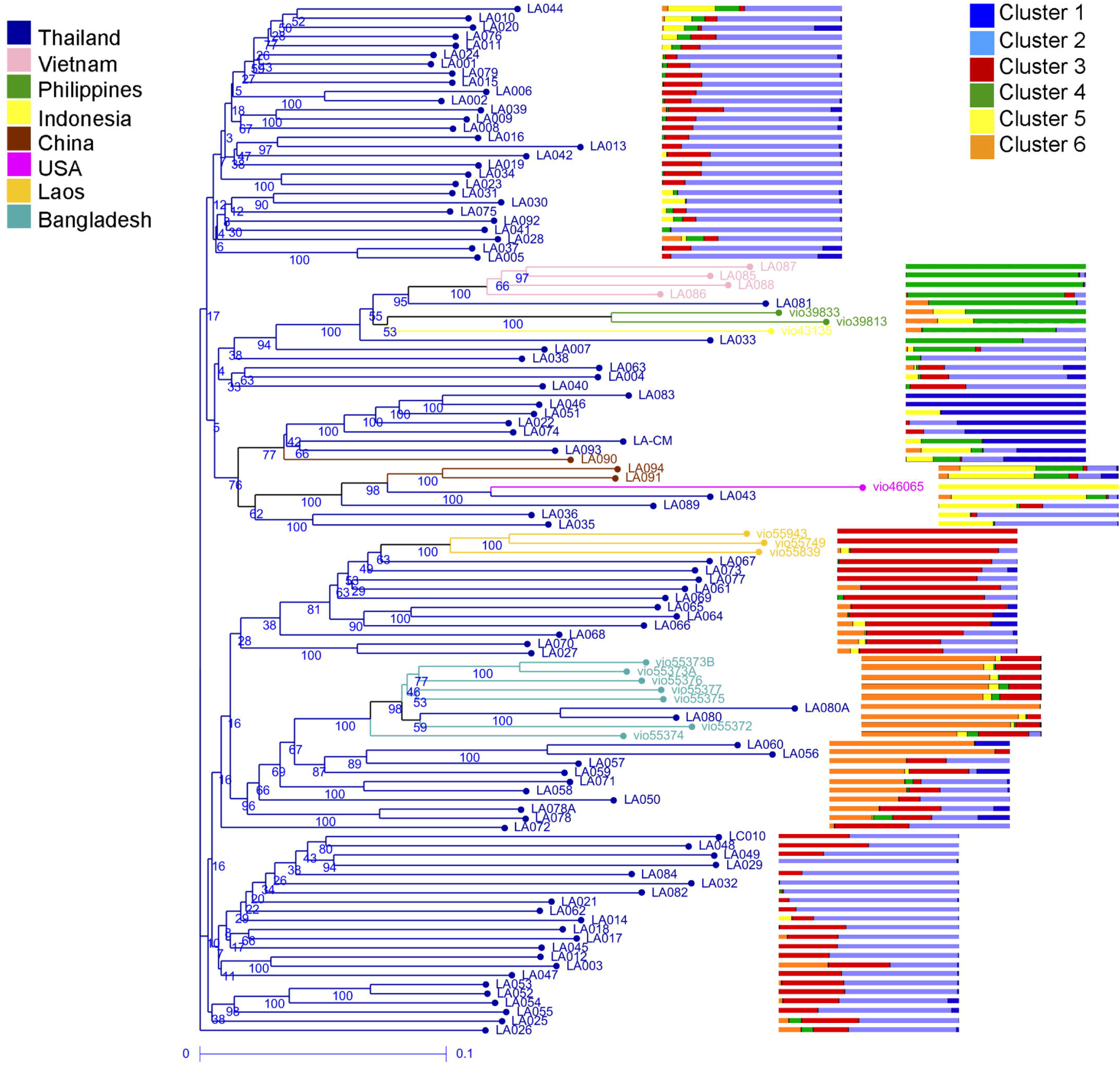
Genetic diversity proportion of 112 *Luffa acutangula* accessions based on country of origin. Branch colors indicate country of origin. Side bars correspond to proportion clustering from Fig. 1

**Fig. 6.**
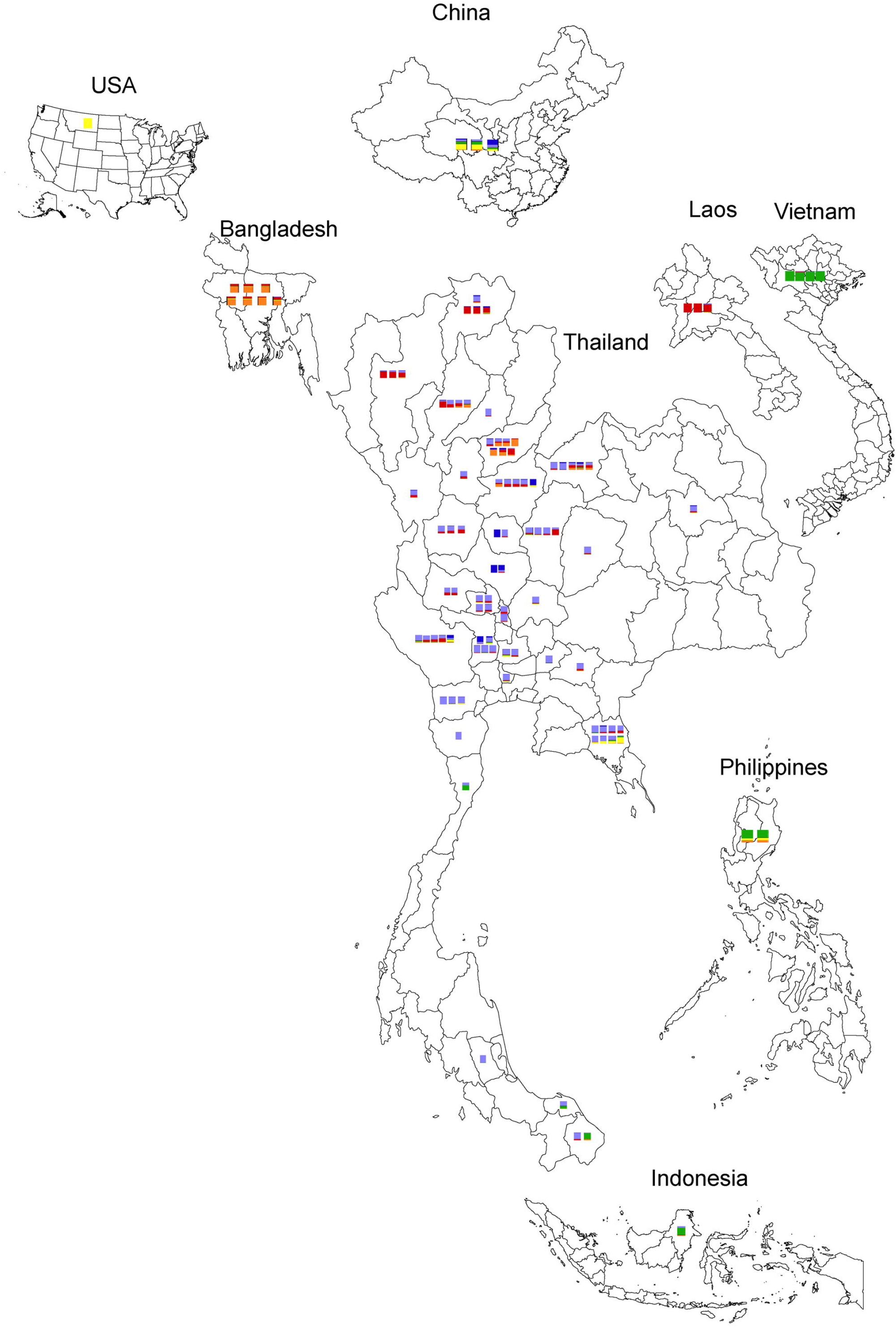
Distribution of *Luffa acutangula* accessions and their corresponding countries. Colors are based on STRUCTURE analysis bars from Fig. 1

### Association of fruit trait and STRUCTURE clustering

The *L. acutangula* germplasm of 112 accessions was diverse for fruit shape and fruit length. Six main types of fruit shapes were used as typical distinguishable phenotypes: elongated slim, elliptical tapered, elongated tapered, elongated elliptical tapered, tapered oblong, and short elliptical tapered (**Fig. 7; Online Resource Online** Resource S2 and **Online** Resource S3). Similar fruit shapes were observed on a previous season of germplasm seed restocking (**Online Resource Online** Resource S7). One-way ANOVA test of fruit length showed highly significant differences between STRUCTURE clusters (p-value = 5.1e-08) (**Online Resource Online** Resource S4). Tukey HSD p-values after adjustment for the multiple comparisons for fruit length showed significant differences for Cluster 1 and Cluster 2, Cluster 1 and Cluster 3, Cluster 1 and 6, Cluster 2 and Cluster 6, and Cluster 4 and Cluster 6 (**Online Resource Online** Resource S5). Other cluster combinations did not show significant differences in fruit length according to Tukey HSD.

**Fig. 7.**
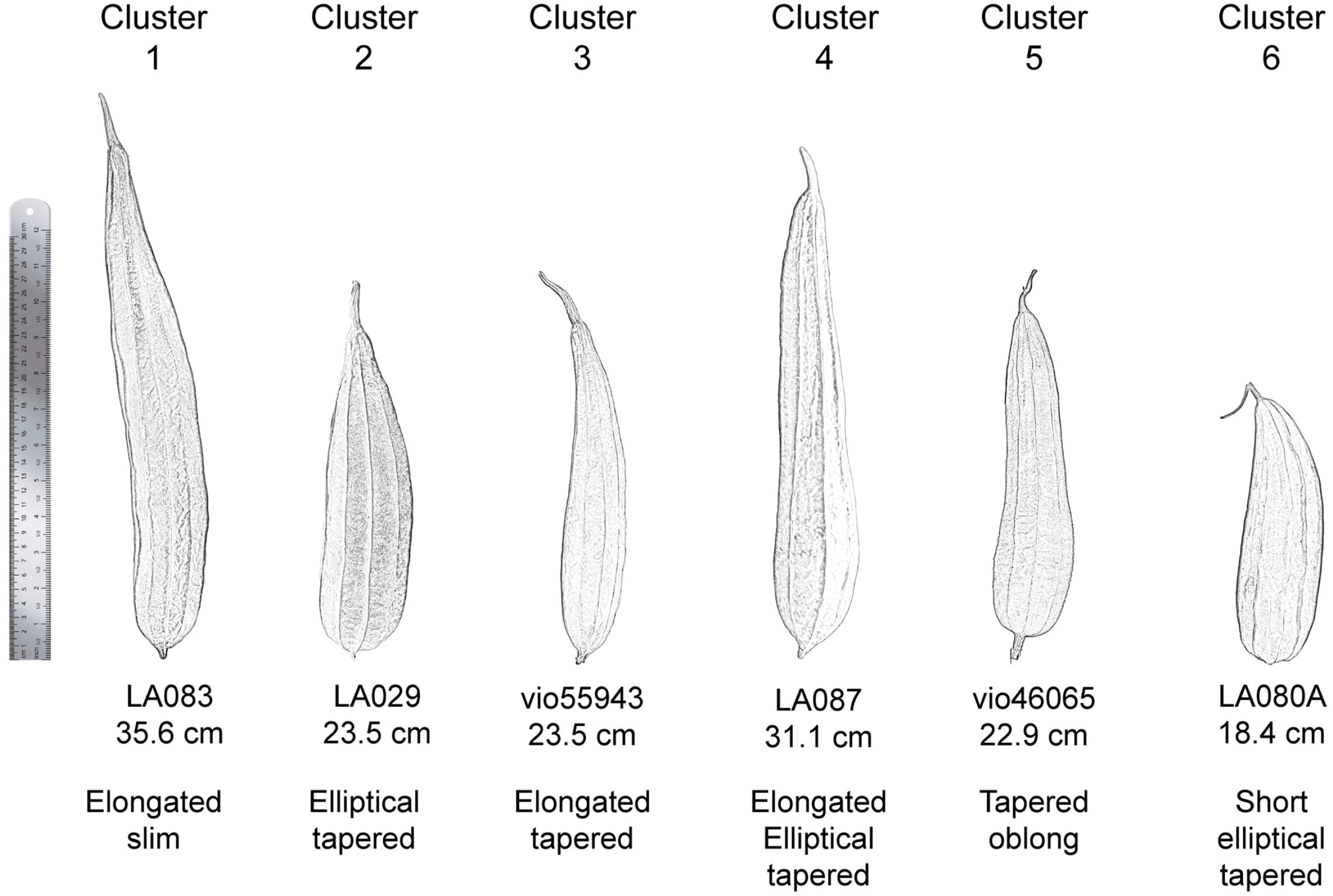
*Luffa acutangula* fruit shape. Each cluster shape is represented by an accession with the least or no admixture (Source: Personal collection credit to Mr. Anucha Wongpraneekul and Grimar A. Perez)

When the fruit shape was defined mainly from the shape of fruits of pure genetic of each subpopulation; fruit shape of subpopulation 1 was elongated slim and showed a long fruit type. Fruit shape of subpopulation 2 was elliptical tapered that had medium length, round blossom end and round stem end. Fruit shape of subpopulation 3 was elongated tapered that had medium to short length, pointed blossom end and pointed stem end. Fruit shape of subpopulation 4 was elongated elliptical tapered with medium to long length with round blossom end and pointed stem end. Subpopulation 5 fruit shape was tapered oblong consisting of medium to short length, oblong shape and a bit round at the bottom end. Fruit shape of subpopulation 6 was short elliptical tapered that are short and quite silmilar to pyriform shape (**Fig. 7**; **Online Resource Online** Resource S6). The fruit shapes of the accessions that showed admixed genetic proportions were affected by the genetic of each subpopulaion representative. This was observed in accessions containing substantial genetic proportions of subpopulation 1, causing a long fruit type or accessions containing genetic of subpopulation 6 which caused a short fruit type.

## Discussion

### Population structure and diversity in *L. acutangula*

In this study, we have been able to demonstrate a consistent six population clustering of the 112 *L. acutangula* accessions. Whole-genome DArTseq based SNPs, geographic area of collection and fruit characteristics were used to study a total of 112 *L. acutangula* accessions. Analysis using STRUCTURE (**Fig. 1**), weighted neighbor-joining method (**Fig. 3**) and PCoA (**Fig. 4**) showed a consistent representation of a total of 6 subpopulations. This grouping consistency was also observed at the level of individual accessions in both STRUCTURE analysis as well as a phylogenetic dendrogram (**Fig. 2**). The subpopulations Cluster 1-6 showed distinct genotypic characteristics that can be put to use in genetic improvement programs.

Geographically, accessions from Thailand made up the majority of Cluster 1 and Cluster 2. Similar correlations between the population structure and geographic origin were observed in *Capsicum* germplasm (Lee et al. 2016). Cluster 1 members were collected from the central plain of Thailand while cluster 2 accessions (n = 62 accessions) were scattered all-over in Thailand. Cluster 3 was composed of accessions from northern Thailand and Laos. This grouping observed in Cluster 3 much emphasizes the similarity in cuisine between the north part of Thailand and Laos. The genotypic characteristics found in the Vietnam accessions (Cluster 4) were found in accessions collected from Southern Thailand. Besides, the genetic proportion of Vietnam accessions found in Cluster 4 was more closely shared with accessions from the Philippines and Indonesia, suggesting a relationship between people from these countries which shared an ancestral history or the possibility that there were immigrations among mainland (Vietnam) and islands in southeast Asia (the Philippines and Indonesia) and also due to the exchange of goods between these countries in the past (Bellina Pryce and Silapanth 2006; Tumonggor et al. 2013). The observed distribution of cluster 4 accessions to the southern part of Thailand point at spreading of seeds along historic commerce ship routes from the Southern Thailand regions to countries such as India and China (Bellina Pryce and Silapanth 2006; Castillo 2011). The primary representative of Cluster 5 genetic was an accession from the USA, which had an almost unmixed nature based on STRUCTURE analysis. Two accessions from China and two accessions from Thailand shared genotypic proportions to this accession in Cluster 5. Cluster 6 was made up of seven accessions from Thailand and seven from Bangladesh, which together was separated into two subgroups in the dendrogram. Cluster 6 genotype characteristic might have been introduced from Nepal region via Bangladesh (Marr et al. 2005), but lack of samples from Myanmar, makes it difficult to confirm the trafficking of this material (Bellina Pryce and Silapanth 2006; Castillo and Fuller 2010).

Cluster 2 and Cluster 6 are two potential groups for breeding programs due to their high diversity when compared across all six subpopulations. Both Cluster 2 and Cluster 6 had high *H_O_* (4.127 and 2.639, respectively) and *H_E_* (0.338 and 0.334, respectively) which indicates a high degree of diversity and substantial heterozygosity. Overall, Cluster 5 displayed the lowest diversity and heterozygosity despite being made up of two accessions from Thailand, two from China and one from the USA.

In a closer look at the genetics of ridge gourd in Thailand such as cluster 1 accessions, we noted that the spreading of their specific SNPs occurred in accessions collected nearby the area where the pure cluster 1 representative accessions had been collected (**Fig. 6**). The same was true for the genetic of cluster 3, which was represented by accessions from Laos. Still, the other accessions in cluster 3 that were domesticated in Thailand were already mixed with the genetics of cluster 2, which is the primary governing genotype of ridge gourd in Thailand.

### Genetic differentiation among populations

Pairwise genetic differentiation (*F*_ST_) was relatively low between Cluster 2 and Cluster 1 (0.089) which is consistent with the PCoA in which the accessions in cluster 1 were found together with the accessions in cluster 2. Cluster 2 and Cluster 3 pairwise genetic differentiation was low (0.086), which also is supported by the PCoA. These two clusters were spread nearby each another, and in the STRUCTURE result, most of the admixed genetics of accessions in Cluster 2 contains the genetics of Cluster 3. Similar observations were seen between Cluster 2 and Cluster 5 (0.147), Cluster 2 and Cluster 6 (0.123), and Cluster 3 and Cluster 6 (0.127) (Table 2). Genetic differentiation of *F*_ST_ lower than 0.05 was defined as low, *F*_ST_ between 0.05 and 0.15 as moderate, *F*_ST_ between 0.15 and 0.25 as high, and *F*_ST_ greater than 0.25 as very high (Frankham et al. 2002; Xu et al. 2019). The low differentiation observed between Cluster 2 with the other four Clusters (Cluster 1, 3, 5 and 6) may reflect the Thailand only composition found in Cluster 2, as well as a substantial diversity found within Cluster 2. Clusters with a pairwise genetic differentiation between 0.150 to 0.210 (Cluster 2 and 4=0.164, Cluster 1 and 3=0.180, Cluster 1 and 6=0.186, Cluster 4 and 6=0.193, Cluster 5 and 6=0.174, Cluster 1 and 5=0.203, Cluster 3 and 5= 0.202, and Cluster 4 and 5=0.210) showed substantial differentiation between populations with minor admixture being exchanged across corresponding paired clusters as seen in the STRUCTURE analysis (**Fig. 1b**) and PCoA. The highest pairwise genetic differentiation occurred between Cluster 1 and Cluster 4 (0.230), and Cluster 3 and Cluster 4 (0.262). These two pairs showed minimal to almost no admixture across each paired cluster as seen in the STRUCTURE analysis, and there was no overlap in the PCoA.

### Thailand germplasm collection and genetic diversity

The within population genetic variation is visible in the phylogenetic tree with corresponding STRUCTURE clustering proportions. Accessions of highly pure genotypes that represent each subpopulation from STRUCTURE analysis are LA083 and LA046 (Cluster 1), LA032 and LA029 (Cluster 2), vio55943 and vio55749 (Cluster 3), LA087 (Cluster 4), vio46065 (Cluster 5), and LA080A (Cluster 6) (**Fig. 1** and **Fig. 5**). The genetic proportions of these accessions can be observed throughout cluster 1 through cluster 6. This observation corresponds to the higher within population diversity found in the AMOVA analysis (Table 3). The AMOVA showed significant population differentiation at the level between subpopulations acquired from STRUCTURE clustering, which support the STRUCTURE assignment of subpopulations. The total population of 112 *L. acutaungula* accessions can be labeled as a panmitic population as most of the observed variance occurred within individuals. The negative AMOVA variance between individuals within populations (−10.59%) indicates no structure in subpopulations, supporting STRUCTURE grouping (Excoffier and Lischer 2009; Misra et al. 2017). This phenomenon is not surprising, given that *L. acutaungula* is an outcrossing species, and it may partly reflect a substantial geneflow between Thailand and countries in Southeast Asia indicating a high genetic exchange between countries. Besides, it might add a new factor to consider in population structure analysis than the usual geopolitical and biological borders (Meirmans 2015).

Part of the South East Asian countries such as Thailand, Laos, Vietnam, Indonesia and Philippines are separated by various biological barriers and marked by geopolitical borders. Despite the proximity of Thailand to Laos and Vietnam, Thailand’s genetic diversity includes a low proportion of genetic characteristics from Vietnam, when compared to the larger proportions from Laos (**Fig. 6**). This observation that may be heavily influenced by cultural similarities when compared to biological barriers or geopolitical borders can also be seen in accessions from Indonesia which to a great extent contain genetic proportions from Vietnam, but none from Laos. However, the Philippines and Vietnam share a similar genetic characteristic in that both of their accessions contain proportions from Vietnam and Bangladesh.

### Association of fruit traits of germplasm collection

Cluster 1 and cluster 4 primarily consisted of long fruits, and elongated slim and elliptical tapered fruit shapes, respectively (**Fig. 7**; **Online Resource Online** Resource S2 and **Online** Resource S3). Cluster 1 and 4 accessions from Thailand, Vietnam, Indonesia, and the Philippines favor long fruits with a wide range of fruit shapes. Cluster 2 was by far the largest subpopulation that was primarily made up of accessions collected in Thailand. Cluster 2 was characterized by medium length fruits with an elliptical tapered fruit shape. However, accessions within Cluster 2 displayed a range of shapes that stemmed from the representative elliptical conical fruit shape. Accessions from Thailand and Laos were grouped in Cluster 3, had medium-length fruits with an elongated tapered fruit shape. This observation between Laos and Thailand represents the similarity in culture, including food shared across both countries. Medium length fruits characterized by elongated elliptical tapered were found in Cluster 5, which consisted of accessions from Thailand, China and the USA. Peculiarly, the accession from the USA in this cluster was perhaps collected from Asia, the reason being that Luffa is native to Asia and not the Americas (Heiser and Schilling 1988; Pessarakli 2016). Cluster 6 was equally composed of accessions from Thailand and Bangladesh. The fruits of Cluster 6 were short in length with a short elliptical tapered fruit shape. This observation in Cluster 6 may indicate a selection of accessions from Bangladesh for breeding purposes in Thailand, or it segregated after being cultivated for some time. The reason for this could be because the majority of accessions in Thailand are widespread in all six subpopulations with a medium to long fruit length. However, fruits from Bangladesh are the only fruits that have significant short fruit lengths found in one out of six subpopulations.

The study found significant genetic diversity in Thailand *L. acutangula* germplasm. Phenotypic information, such as the fruit length and fruit shape, positively followed suit in the variation observed across the six populations inferred by the genotypic data. At the same time, geographical provenance was reflected in the clustering analysis, which also showed a relationship to the phenotypic information. This ample diversity serves as a strong catalyst to utilize such germplasm for breeding programs which can be geared towards farmer and consumer preference.

## Supporting information

Online Resource

## Acknowledgements

Special thanks to the Tropical Vegetable Research Center and the World Vegetable Center for providing seeds, and to Mr. Anucha Wongpraneekul and Ms. Waraporn Sinsathapornpong for their help with field management of ridge gourd and photos of the fruits.

## Supplementary information (SI)

**Online Resource S1.** Passport Data of *L. acutangula* accessions. *TVRC* Tropical Vegetable Research Center, *AVRDC* World Vegetable Center.

**Online Resource S2.** Descriptive statistics of *Luffa acutangula* fruit length (cm).

**Online Resource S3.** Box plots of fruit length (cm) by STRUCTURE clustering.

**Online Resource S4.** One-way ANOVA based on STRUCTURE grouping. ***P-value < 0.001.

**Online Resource S5.** Tukey HSD based on STRUCTURE grouping with a 95% family-wise confidence level. *Diff* mean difference between two groups, *Lwr* lower endpoint of the interval, *Upr* upper endpoint of the interval, *P adj* p-value after adjustment for the multiple comparisons.

**Online Resource S6.** Fruit shape of *Luffa acutangula* accessions based on STRUCTURE grouping.

**Online Resource S7.** Fruit shape of *Luffa acuntagula* representatives based on STRUCTURE grouping.

## References

Barilli E, Cobos MJ, Carrillo E, Kilian A, Carling J, Rubiales D (2018) A high-density integrated DArTseq SNP-based genetic map of *Pisum fulvum* and identification of QTLs controlling rust resistance. Frontiers in Plant Science 9:167

Bellina Pryce B, Silapanth P (2006) Weaving cultural identities on trans-Asiatic networks: Upper Thai-Malay Peninsula–an early socio-political landscape. Bulletin de l’École française d’Extrême-Orient:257–293

Bretting P, Widrlechner MP (1995) Genetic markers and plant genetic resource management.

Bruno WJ, Socci ND, Halpern AL (2000) Weighted neighbor joining: a likelihood-based approach to distance-based phylogeny reconstruction. Molecular biology and evolution 17:189–197

Castillo C (2011) Rice in Thailand: the archaeobotanical contribution. Rice 4:114

Castillo C, Fuller DQ (2010) Still too fragmentary and dependent upon chance? Advances in the study of early Southeast Asian archaeobotany. Fifty years of archaeology in Southeast Asia: Essays in honour of Ian Glover:91–111

Chesnokov YV, Artemyeva A (2015) Evaluation of the measure of polymorphism information of genetic diversity. Сельскохозяйственная биология

Doyle JJ, Doyle JL (1987) A rapid DNA isolation procedure for small quantities of fresh leaf tissue. Phytochemical Bulletin 19:11–15

Dray S, Dufour AB (2007) The ade4 package: implementing the duality diagram for ecologists. Journal of statistical software 22:1–20

Evanno G, Regnaut S, Goudet J (2005) Detecting the number of clusters of individuals using the software STRUCTURE: a simulation study. Molecular ecology 14:2611–2620

Excoffier L, Lischer H (2009) Arlequin 3.5. An Integrated Software Package for Population Genetics Data Analysis Computational and Molecular Population Genetics Lab (CMPG) Institute of Ecology and Evolution University of Berne, Switzerland

Filipowicz N, Schaefer H, Renner SS (2014) Revisiting *Luffa* (Cucurbitaceae) 25 years after C. Heiser: species boundaries and application of names tested with plastid and nuclear DNA sequences. Systematic Botany 39:205–215

Francis RM (2017) pophelper: an R package and web app to analyse and visualize population structure. Molecular ecology resources 17:27–32

Frankham R, Briscoe DA, Ballou JD (2002) Introduction to conservation genetics. Cambridge university press

Heiser CB, Schilling EE (1988) Phylogeny and distribution of *Luffa* (Cucurbitaceae). Biotropica:185–191

Holsinger DAFKE (1991) Genetics and conservation of rare plants. Oxford University Press on Demand

Hoque S, Rabbani M (2009) Assessment of genetic relationship among landraces of Bangladeshi ridge gourd (*Luffa acutangula* Roxb.) using RAPD markers. Journal of Scientific Research 1:615–623

Jombart T, Ahmed I (2011) adegenet 1.3-1: new tools for the analysis of genome-wide SNP data. Bioinformatics 27:3070–3071

Joshi BK, KC HB, Tiwari RK, Ghale M, Sthapit BR (2004) Descriptors for sponge gourd (*Luffa cylindrica* (L.) Roem.).

Kamvar ZN, Tabima JF, Grünwald NJ (2014) Poppr: an R package for genetic analysis of populations with clonal, partially clonal, and/or sexual reproduction. PeerJ 2:e281

Katewa S, Chaudhary B, Jain A (2004) Folk herbal medicines from tribal area of Rajasthan, India. Journal of Ethnopharmacology 92:41–46

Kilian A, Wenzl P, Huttner E, Carling J, Xia L, Blois H, Caig V, Heller-Uszynska K, Jaccoud D, Hopper C (2012) Diversity arrays technology: a generic genome profiling technology on open platforms. Data production and analysis in population genomics. Springer, pp 67–89

Lee H-Y, Ro N-Y, Jeong HJ, Kwon JK, Jo J, Ha Y, Jung A, Han JW, Venkatesh J, Kang B-C (2016) Genetic diversity and population structure analysis to construct a core collection from a large *Capsicum* germplasm. BMC genetics 17:142

Leffler EM, Bullaughey K, Matute DR, Meyer WK, Segurel L, Venkat A, Andolfatto P, Przeworski M (2012) Revisiting an old riddle: what determines genetic diversity levels within species? PLoS Biol 10:e1001388

Mardis ER (2008) The impact of next-generation sequencing technology on genetics. Trends in genetics 24:133–141

Marr KL, Bhattarai NK, Xia Y-M (2005) Allozymic, morphological, and phenological diversity in cultivated *Luffa acutangula* (Cucurbitaceae) from China, Laos, and Nepal, and Allozyme Divergence between *L. acutangula* and *L. aegyptiaca*. Economic Botany 59:154–165

Meirmans PG (2015) Seven common mistakes in population genetics and how to avoid them. Molecular ecology 24:3223–3231

Misra S, Srivastava AK, Verma S, Pandey S, Bargali SS, Rana TS, Nair KN (2017) Phenetic and genetic diversity in Indian *Luffa* (Cucurbitaceae) inferred from morphometric, ISSR and DAMD markers. Genetic resources and crop evolution 64:995–1010

Morton JF (1981) Atlas of medicinal plants of middle America: Bahamas to Yucatan. [Volume 1]. Thomas

Pandey S, Ansari W, Choudhary B, Pandey M, Jena S, Singh A, Dubey R, Singh B (2018) Microsatellite analysis of genetic diversity and population structure of hermaphrodite ridge gourd (*Luffa hermaphrodita*). 3 Biotech 8:17

Perrier X, Jacquemoud-Collet JP (2006) DARwin software.

Pessarakli M (2016) Handbook of Cucurbits: growth, cultural practices, and physiology. CRC Press

Prakash K, Pandey A, Radhamani J, Bisht I (2013) Morphological variability in cultivated and wild species of *Luffa* (Cucurbitaceae) from India. Genetic resources and crop evolution 60:2319–2329

Prakash K, Pati K, Arya L, Pandey A, Verma M, Prakash K, Pati K, Arya L, Pandey A, Verma M (2014) Population structure and diversity in cultivated and wild *Luffa* species. Biochemical Systematics and Ecology:165–170

Pritchard JK, Stephens M, Donnelly P (2000) Inference of population structure using multilocus genotype data. Genetics 155:945–959

Raman H, Raman R, Kilian A, Detering F, Carling J, Coombes N, Diffey S, Kadkol G, Edwards D, McCully M (2014) Genome-wide delineation of natural variation for pod shatter resistance in *Brassica napus*. LoS One 9:e101673

Samvatsar S, Diwanji V (2000) Plant sources for the treatment of jaundice in the tribals of Western Madhya Pradesh of India. Journal of Ethnopharmacology 73:313–316

Sastri B (1962) The wealth of India, vol. 6. Council of Scientific and Industrial Research, New Delhi:382–383

Soladoye MO, Adebisi AA (2004) *Luffa acutangula* (L.) Roxb. In: Grubben GJH, Denton OA (eds)

Tumonggor MK, Karafet TM, Hallmark B, Lansing JS, Sudoyo H, Hammer MF, Cox MP (2013) The Indonesian archipelago: an ancient genetic highway linking Asia and the Pacific. Journal of human genetics 58:165–173

Von Mark VC, Kilian A, Dierig DA (2013) Development of DArT marker platforms and genetic diversity assessment of the US collection of the new oilseed crop lesquerella and related species. PLoS One 8:e64062

Xu Y, Mai JW, Yu BI, Hu HX, Yuan L, Jashenko R, Ji R (2019) Study on the genetic differentiation of geographic populations of *Calliptamus italicus* (Orthoptera: Acrididae) in sino-kazakh border areas based on mitochondrial COI and COII genes. Journal of economic entomology 112:1912–1919

Zhang S, Hu J, Xu S (2008) Developmental genetic analysis of fruit shape traits under different environmental conditions in sponge gourd (*Luffa cylindrical* (L) Roem. Violales, Cucurbitaceae). Genetics and Molecular Biology 31:708–710

