## Supplementary figures and images for "Genetic diversity and population structure of *Luffa acutangula* accessions in a Thailand collection using SNP markers"

### Online Resource S3.tif

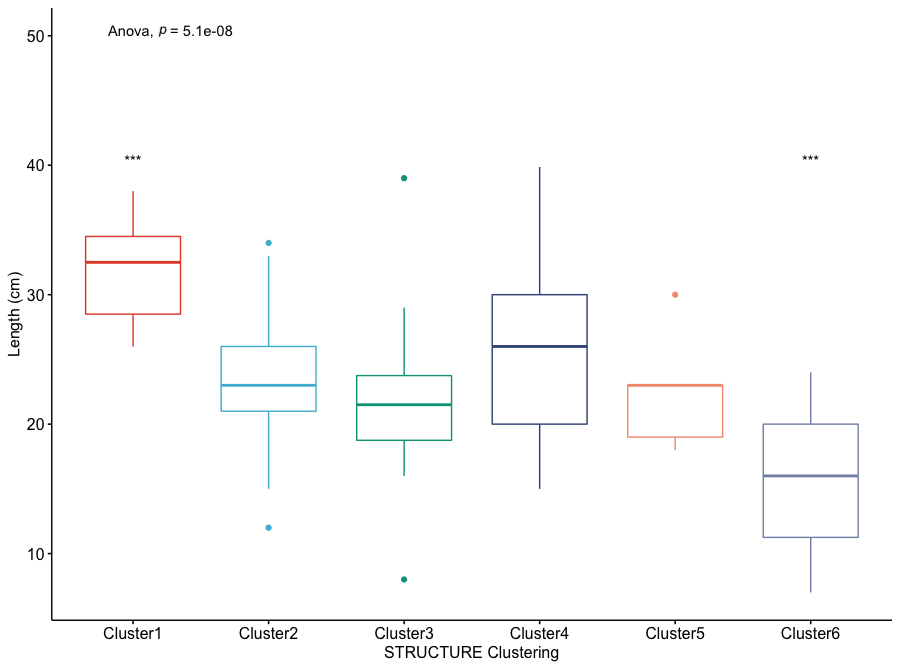

### Online Resource S6.tif

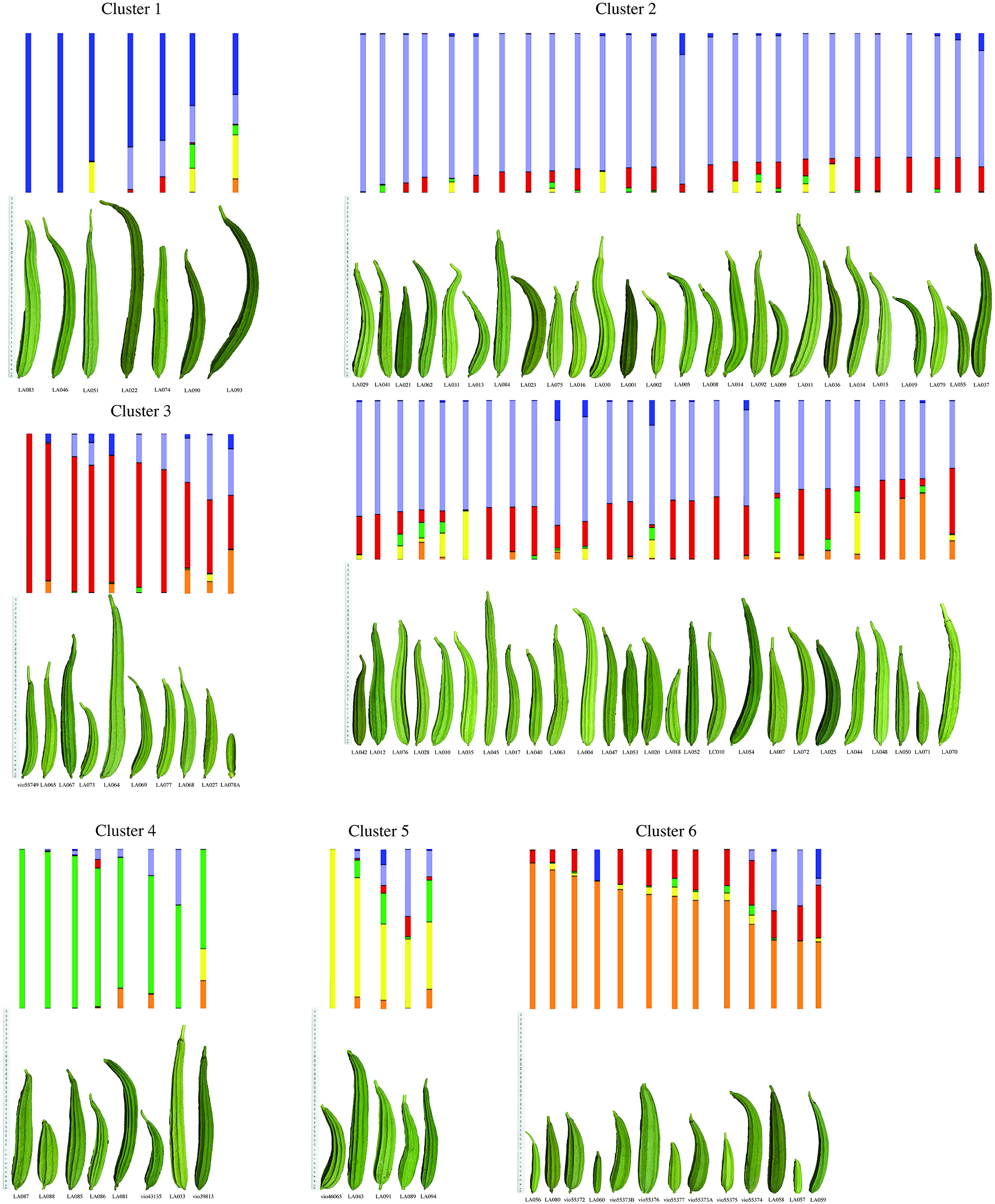

### Online Resource S7.tif

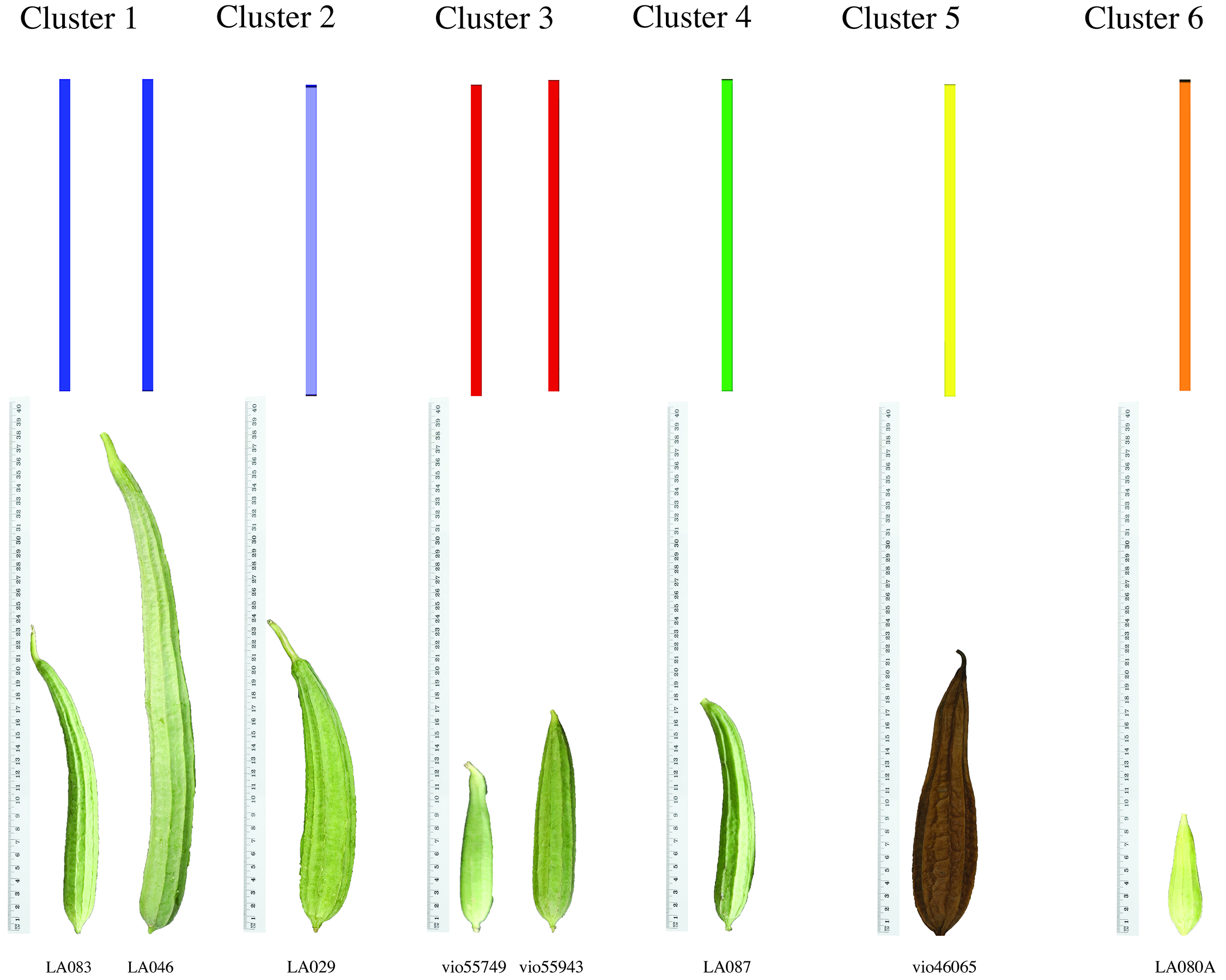
